# The endosymbiont of *Epithemia clementina* is specialized for nitrogen fixation within a photosynthetic eukaryote

**DOI:** 10.1101/2023.03.08.531752

**Authors:** Solène L.Y. Moulin, Sarah Frail, Jon Doenier, Thomas Braukmann, Ellen Yeh

**Affiliations:** Department of Pathology, Stanford School of Medicine, Stanford, California, USA; Department of Biochemistry, Stanford School of Medicine, Stanford, California, USA; Department of Microbiology & Immunology, Stanford School of Medicine, Stanford, California, USA; Chan Zuckerberg Biohub – San Francisco, San Francisco, CA 94158

**Author notes:** 299 Campus Dr, Fairchild 317, Stanford, CA 94305. Author Contributions: S.L.Y.M., and E.Y. designed research; S.L.Y.M., S.F., and T.B. performed research; S.L.Y.M., S.F., T.B. and J.D analyzed data; and S.L.Y.M., and E.Y. wrote the paper. Competing Interest Statement: The authors declare no competing interest. Classification: Major: Biological sciences; Minor: cell biology.

**Keywords:** Endosymbiosis, Nitrogen fixation, cyanobacteria, diatom.

## Abstract

*Epithemia* spp. diatoms contain obligate, nitrogen-fixing endosymbionts, or “diazoplasts”, derived from cyanobacteria. These algae are a rare example of photosynthetic eukaryotes that have successfully coupled oxygenic photosynthesis with oxygen-sensitive nitrogenase activity. Here, we report a newly-isolated species, *E. clementina*, as a model to investigate endosymbiotic acquisition of nitrogen fixation. To detect the metabolic changes associated with endosymbiotic specialization, we compared nitrogen fixation, associated carbon and nitrogen metabolism, and their regulatory pathways in the *Epithemia* diazoplast with its close, free-living cyanobacterial relative, *Crocosphaera subtropica*. Unlike *C. subtropica*, we show that nitrogenase activity in the diazoplast is concurrent with, and even dependent on, host photosynthesis and no longer associated with cyanobacterial glycogen storage suggesting carbohydrates are imported from the host diatom. Carbohydrate catabolism in the diazoplast indicates that the oxidative pentose pathway and oxidative phosphorylation, in concert, generates reducing equivalents and ATP and consumes oxygen to support nitrogenase activity. In contrast to expanded nitrogenase activity, the diazoplast has diminished ability to utilize alternative nitrogen sources. Upon ammonium repletion, negative feedback regulation of nitrogen fixation was conserved, however ammonia assimilation showed paradoxical responses in the diazoplast compared with *C. subtropica*. The altered nitrogen regulation likely favors nitrogen transfer to the host. Our results suggest that the diazoplast is specialized for endosymbiotic nitrogen fixation. Altogether, we establish a new model for studying endosymbiosis, perform the first functional characterization of this diazotroph endosymbiosis, and identify metabolic adaptations for endosymbiotic acquisition of a critical biological function.

## Significance Statement

Nitrogen is a limiting nutrient for photosynthetic productivity in natural ecosystems and especially in agriculture. Abundant nitrogen in the atmosphere must be fixed into bioavailable ammonia, a reaction performed in bacteria that is exquisitely sensitive to oxygen produced during photosynthesis. How then can plants and other photosynthetic eukaryotes acquire nitrogen fixation function? We investigate a photosynthetic alga that hosts nitrogen-fixing cyanobacteria as endosymbionts residing inside the eukaryotic cell. Our results reveal critical metabolic adaptations associated with the transition from a free-living cyanobacteria to a nitrogen-fixing specialist endosymbiont inside a photosynthetic host cell. Elucidating how photosynthesis and nitrogen fixation are coupled in this eukaryotic cell holds important lessons for how to bioengineer nitrogen-fixing crop plants towards the goal of sustainable agriculture.

## Introduction

Nitrogen composes 78% of the atmosphere; however, it is present in the form of dinitrogen (N_2_), an inert gas, which cannot be directly incorporated into organic molecules such as DNA or protein. Nitrogen fixation, a critical biological reaction, is the reduction of N_2_ into bioavailable ammonia catalyzed by the enzyme nitrogenase. Biological nitrogen fixation is largely restricted to certain genera of bacteria. Nitrogen availability determines photosynthetic productivity in both natural environments and agriculture, such that the activity of nitrogen-fixing organisms, or diazotrophs, is a critical source of bioavailable nitrogen for plants and other photosynthetic organisms. To bypass the dependence of food production on biological nitrogen fixation, modern agriculture uses energy from fossil fuels to chemically fix nitrogen and produce nitrogen-rich fertilizer, which is estimated to feed 45% of the world population (1). However, synthetic fertilizers have significant negative environmental consequences including greenhouse gas emissions, soil degradation, and coastal dead zones (eutrophication) with runoff nitrogen polluting waterways (2–4).

While a nitrogen-fixing plant is an obvious solution, the closest example is legumes which host nitrogen-fixing symbionts in root nodules. Attempts to introduce nitrogen fixation into plant cells by expressing nitrogenase genes have met significant challenges (5, 6). On one hand, nitrogenase requires 16 ATP and 8 NADPH molecules to fix a single N_2_ molecule, energy that can be supplied by the photosynthetic electron transfer chain. Thus, coupling nitrogen fixation and photosynthesis in a single cell is energetically advantageous. On the other hand, however, nitrogenase is extremely sensitive to irreversible inactivation by oxygen which is produced during photosynthesis, implying that photosynthesis and nitrogen fixation are biochemically incompatible. Instead, throughout evolution, diverse photosynthetic eukaryotes have coupled these two biologically essential reactions via symbiosis with nitrogen-fixing bacteria (7).

Amongst these diverse symbiotic interactions, a few rare microalgae have acquired diazotrophic cyanobacterial endosymbionts allowing them to grow in nitrogen-limited environments. *Crocosphaera* spp. cyanobacteria (previously named *Cyanothece*) appear to be a privileged endosymbiotic partner (8). Several microalgae, the diatoms *Epithemia* spp. and *Climacodium frauenfeldianum* (9) as well as the coccolithophore *Braarudosphaera bigelowii* (10), have permanent endosymbionts closely related to the *Crocosphaera* genus, specifically *Crocosphaera subtropica* (*Cyanothece* sp. ATCC51142). Within their photosynthetic hosts, most of these cyanobacterial endosymbionts have lost their photosynthetic function and become specialized for nitrogen fixation. The remarkable success of *Crocosphaera* cyanobacteria to transition from free-living diazotrophic phototroph to endosymbiotic diazotrophic heterotroph in multiple eukaryotic lineages suggests pre-existing traits that predispose them to endosymbiotic acquisition. For example, *C. subtropica* can grow mixotrophically (i.e., relying on photosynthesis to fix carbon or using carbon substrates from the environment), which may favor its success as an endosymbiotic partner (11, 12).

*Crocosphaera*’s partnership with *Epithemia* diatoms has been particularly successful. Diverse *Epithemia* species containing endosymbionts are widespread in freshwater habitats globally and have recently been found in marine environments (13). Their remarkable ability to fix both nitrogen and carbon plays a critical ecological role in the food web of aquatic environments (14, 15). The endosymbionts, first described as “spheroid bodies” within *Epithemia gibba* (formerly *Rhopalodia gibba* (16)), were shown to encode an intact *nif* gene cluster and demonstrated nitrogen fixation activity enabling the host diatom to grow in nitrogen-depleted environments (17–20). Hereafter, we will refer to these nitrogen-fixing endosymbionts as diazoplast, combining “diazo-” meaning ‘relating to the group N_2_’ and “-plast” from the Greek *plastos* meaning ‘formed’. Sequencing of diazoplast genomes from two *Epithemia* species revealed significant genome reduction, including loss of most of the genes involved in photosynthesis, implying that diazoplasts are dependent on the diatom for their metabolic needs (20–22). Consistent with their being obligate endosymbionts, diazoplasts have never been observed outside their diatom, and isolated diazoplasts cannot be grown in laboratory cultures outside the host cells. Diazoplasts are not only vertically transmitted during cell division but show uniparental inheritance during sexual reproduction, demonstrating robust mechanisms of inheritance (23). Although *Epithemia* spp. are morphologically diverse, their diazoplasts are believed to derive from a single endosymbiotic event with a cyanobacteria from *Crocosphaera* genus (24). Dated at approximately 12 Mya (25), this is the most recent obligate primary endosymbiosis documented to date. Finally, several *Epithemia* strains isolated from environmental samples have been successfully grown in laboratory cultures (26). Overall, the diazoplasts of *Epithemia* diatoms are an ideal case study both for the acquisition of nitrogen fixation function by a photosynthetic eukaryote and for investigating evolution associated with organellogenesis (21).

Though *Epithemia*’s diazoplasts were described nearly half a century ago, little is known about this functionally unique and ecologically widespread endosymbiotic interaction. So far, the most detailed molecular studies have focused on the extensive genome reduction in the diazoplast suggestive of unprecedented metabolic dependence on the host cell (20, 21). While the pattern of reduction and retention in the diazoplast genome suggests new cell biology, such as the energetic coupling of diatom photosynthesis to diazoplast nitrogen fixation, functional studies to support these hypotheses have been limited. Herein, we systematically compare nitrogen fixation and nitrogen assimilation, as well as their regulatory pathways in the *Epithemia* diazoplast with a close free-living relative. The diazoplast was investigated in a newly isolated *Epithemia* species, *E. clementina*, while *C. subtropica* served as a proxy for the ancestral free-living cyanobacteria. Our studies reveal key adaptations accompanying the transition from free-living to endosymbiont that allowed diazoplasts to become specialized for nitrogen fixation in a photosynthetic eukaryote.

## Results

### *Epithemia clementina* is a new model system to study evolution of diazoplasts

*Epithemia* diatoms are widely distributed in freshwater environments globally, and laboratory cultures have previously been established from isolates collected in the US, Germany, and Japan. Of these cultures, only one *Epithemia* strain has been deposited in a culture collection (*Rhopalodia gibba*, FD213, UTEX); however, we were unable to establish cultures from this isolate. Previous study has reported *Epithemia* sp. in the Eel river in Northern California (14). Therefore, to obtain a strain of *Epithemia* for laboratory studies, we collected water samples from a freshwater stream near Butano State Park, CA. The samples were serially diluted over a period of 8 weeks in nitrogen and carbon-free media to isolate diazotrophic phototrophs. Most isolates were dominated by green-colored filamentous cyanobacteria, likely *Nostoc* spp. that are ubiquitous phototrophic diazotrophs. However, a few isolates were visibly brown and consisted of diatoms resembling previously described *Epithemia* spp. that contained endosymbionts. Consistent with their unique ability to fix nitrogen, these diatoms were the only eukaryotic algae that were isolated from passage in nitrogen-free media.

Under light microscopy, the isolated alga was approximately 20 µm long and resembled a wedge of an orange (Fig. 1*A*). Because morphologic and phylogenetic characterizations (detailed below) suggest it is a new species, it was named *Epithemia clementina*. Like other members of its genus, it is a raphid pennate diatom possessing lunate, planar valve faces and prominent internal costae. It has an asymmetrical growth of its gridle bands and shows the presence of marginal raphes on its concave side. Morphologically, the shape and pattern associated with the valve does not match other described *Epithemia* species (Fig. S1*A*). Similar to other *Epithemia* diatoms, *E. clementina* contains endosymbionts appearing as spheroid bodies within the diatom both under light microscopy and by fluorescence microscopy after staining for nucleic acids (Fig. 1*A* and *B*). Nucleic acid staining also revealed the presence of bacteria attached to the frustule of *E. clementina.* Consistent with previous characterization of diazoplasts, the endosymbionts were colorless in light microscopy and did not have any signal in fluorescence microscopy, indicating a lack of photosynthetic pigments (Fig. 1*C* and *D*, Fig. S1*B*).

**Figure 1.**
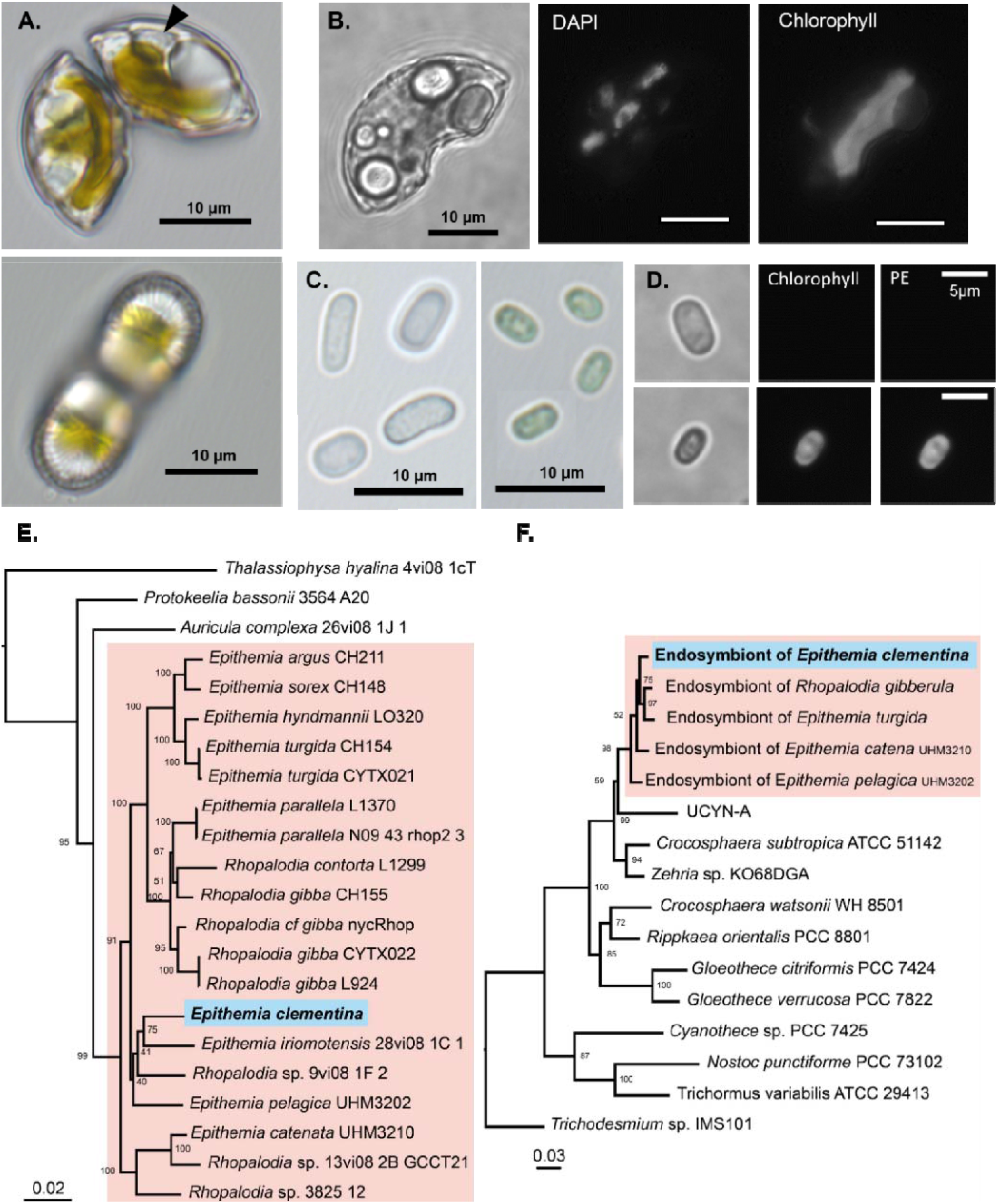
Identification of *Epithemia clementina*. (A) Light microscopy of *Epithemia clementina* apex (top) and valve/ventral region (bottom) with “spheroid bodies” (black arrow). (B) Epifluorescence microscopy of *E. clementina* comparing brightfield (left), DAPI stain (middle), and chlorophyll florescence (right). (C) Light microscopy of isolated endosymbionts released from crushed *E. clementina* (left), compared with free-living *C. subtropica* (right). (D) Epifluorescence microscopy of isolated endosymbionts (top) and *C. subptropica* (bottom) comparing brightfield (left), chlorophyll (Chl) fluorescence (middle), and phycoerythrin (PE) fluorescence (right). (E) Multigene phylogeny of *E. clementina* and related diatoms based on the ribosomal small subunit RNA (18S rRNA), psbC, and rbcL genes. (F) Multigene phylogeny of the *E. clementina* endosymbionts and related cyanobacteria, based on the ribosomal small subunit RNA (16S rRNA) and *nifH* genes. Phylogeny is supported by bootstrap values, and phylogeny scales are in units of nucleotide substitutions per site. Accession numbers for all sequences are provided in Source Data file (Table S4).

The morphologic classification was supported by molecular phylogeny. Phylogenetic analysis based on a selection of conserved genes from diatoms confirmed that this undescribed species is a member of the *Epithemia* genus, most closely related to *Epithemia iriomotensisa* (Fig. 1*E*) (16). To identify the origin of the endosymbiont, we also performed a phylogenetic analysis of conserved cyanobacterial genes which showed that the endosymbiont of *E. clementina* endosymbiont is of the same origin as endosymbionts previously found in *Epithemia* spp. (Fig. 1*F*) (24). The diazoplast genome was assembled with a hybrid assembly strategy using Nanopore long-read and Illumina short-read sequencing of a metagenomic sample of our lab isolate containing *E. clementina* and associated bacteria. The genome confirmed that the endosymbiont was indeed a diazoplast, containing an intact *nif* cluster as observed in other *Epithemiaceae* (20, 21). Having observed bacteria closely associated with *E. clementina* in the monoalgal cultures that could not be removed by dilution, detergent, or antibiotic treatment, we took advantage of our metagenomic sequencing data to characterize the bacterial community of *E. clementina*. 62 metagenome-assembled genomes (MAGs) from taxonomically diverse bacteria were obtained in addition to the diazoplast genome (Fig. S2, Table S1). None of the bacteria were assigned as phototrophs, though the presence of *nif* genes was detected in 2 MAGs related to *Methyloversatilis* sp. RAC08 (∼90% identity) and *Hyphomicrobium* sp. DMF-1 (∼75-80% identity). Based on sequencing depth, the diazoplast was the most abundant bacterial genome and accounted for 19.2% of bacterial genomes detected, compared to <0.3% for the two nif-containing MAGs, suggesting that it is the primary source of nitrogenase activity in our cultures.

### The diazoplast has lost temporal regulation of nitrogenase

Having established laboratory cultures of *E. clementina* to study its diazoplast, we also obtained cultures of *C. subtropica,* which is closely related to the diazoplast (Fig. 1*F*), to serve as a proxy for the free-living ancestor of the diazoplast. We first compared the temporal regulation of nitrogen fixation activity in free-living *C. subtropica* and *E. clementina*’s diazoplast. Nitrogenase is sensitive to deactivation by oxygen; as a result, mechanisms to separate nitrogenase activity and oxygenic photosynthesis either spatially or temporally have evolved in cyanobacteria like *C. subtropica* that perform both photosynthesis and nitrogen fixation. Studies of *C. subtropica* demonstrated tight transcriptional and post-translational regulation of nitrogenase, such that nitrogen fixation only occurs at night when photosynthesis is inactive (27–30). Using an acetylene reduction assay (ARA), we confirmed that the nitrogenase activity of *C. subtropica* was restricted to nighttime with a strong diel pattern that peaked around 4 hours into the night (Fig. 2*A*) (29, 31, 32).

**Figure 2.**
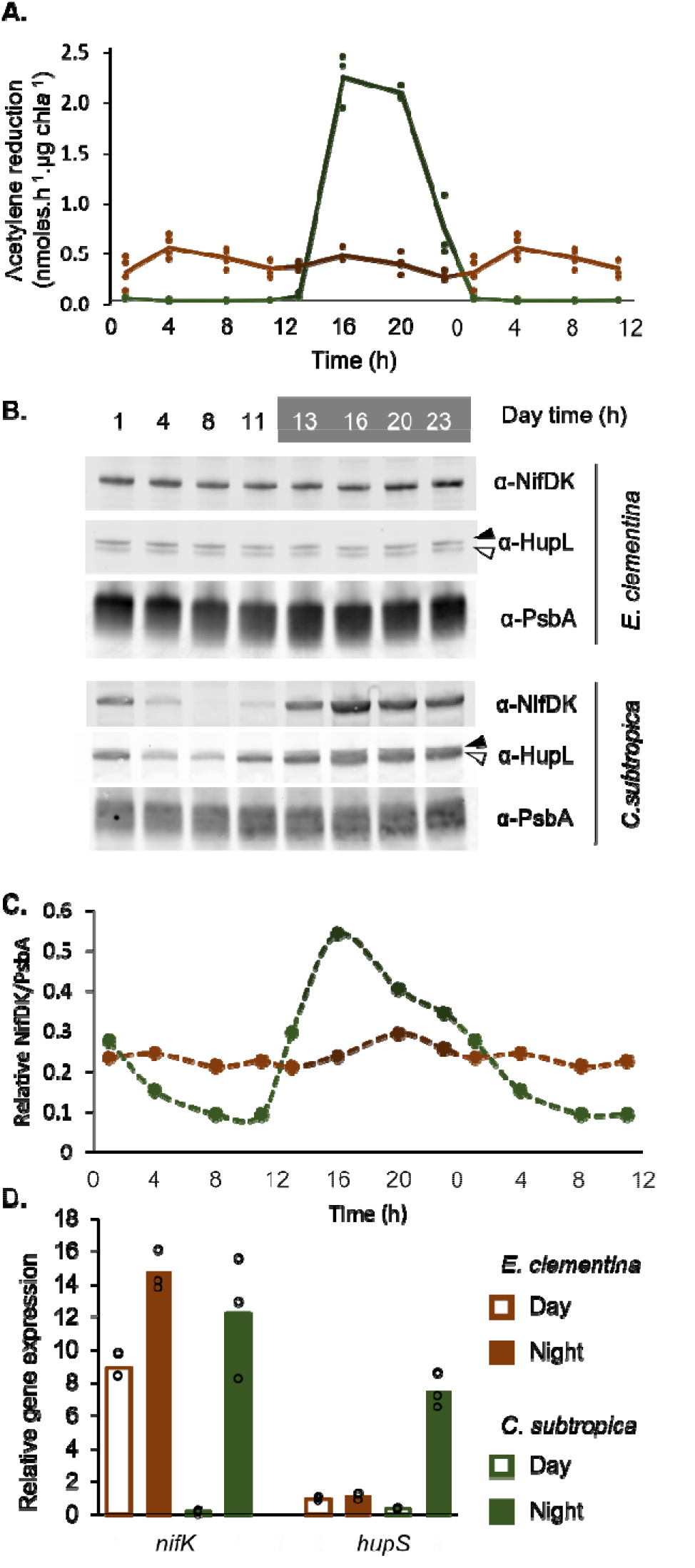
Diel pattern of nitrogen fixation in *E. clementina* and *C. subtropica* ATCC 51142. (A) Nitrogenase activity assessed by acetylene reduction assay over a diel cycle in *E. clementina* (brown) and *C. subtropica* ATCC 51142 (green) normalized by chlorophyll content. Timepoints (1, 4, 8 and 11h) were duplicated for better visualization of the cycle. (B) Corresponding immunoblot for nitrogenase (NifDK) and hydrogenase uptake (HupL). The black arrow points to the apo-HupL and the white arrow points to the mature HupL protien. (C) Quantification of NifDK to PsbA ratio from B. Timepoints (1, 4, 8 and 11h) were duplicated for better visualization of the cycle. (D) Relative gene expression of *nifK* and *hupS* in *E. clementina* (brown) and *C. subtropica* (green) during early day (open bars) and early night (solid bars). Mean values are plotted with individual biological replicates indicated by data points.

In contrast to *C. subtropica,* the diazoplast is no longer photosynthetic, thus we assessed if the spatial separation from photosynthesis performed by the host, has affected the diel regulation of the nitrogenase activity in the diazoplast. Studies of other *Epithemia* spp. have demonstrated daytime nitrogenase activity concurrent with photosynthesis, indicating a remodeling of the temporal regulation (13, 17, 18). Consistent with these previous reports and in contrast to free-living *C. subtropica*, *E. clementina* showed nitrogenase activity throughout the day-night cycle with no clear diel pattern (Fig. 2*A*). In *C. subtropica,* immunoblotting for the NifDK subunits of nitrogenase showed a diel pattern matching the nitrogenase activity, while in *E. clementina*, NifDK proteins abundance was stable throughout the day-night cycle (Fig. 2*B* and *C*). We also assessed the abundance of HupL, the large subunit of the uptake hydrogenase which recycles hydrogen produced by nitrogenase and is often co-expressed with *nif* genes (33). HupL protein also presented a diel variation being abundant at night and fading out during the day in *C. subtropica*, while HupL protein abundance was stable in the diazoplast (Fig. 2*B*). Finally, we performed quantitative RT-qPCR to detect transcript levels of the NifK subunit of nitrogenase and HupS, the small subunit of the uptake hydrogenase. Both *nifK* and *hupS* showed tightly-regulated nighttime expression in *C. subtropica* while the diazoplast showed constitutive *nifK* and *hupS* expression (Fig. 2*D*). Taken together, the activity, protein abundance, and transcript levels of nitrogenase demonstrate that the global temporal regulation of nitrogen fixation in free-living *C. subtropica* is lost in the diazoplast of *E. clementina*.

### Diazoplast nitrogen fixation is coupled to host photosynthesis and carbohydrate storage

Biological nitrogen fixation is an energy-intensive reaction; therefore, the metabolism of diazotrophs must have the ability to supply large quantities of both ATP and reducing equivalents to the nitrogenase. In free-living *Crocosphaera*, the energy for nitrogen fixation is provided by photosynthesis, which converts sunlight into chemical energy in the form of glucose. However, because photosynthesis is segregated temporally from nitrogen fixation in *Crocosphaera*, glucose produced during the day is stored in glycogen granules and catabolized at night when nitrogenase is active (29, 32). A global transcriptomic analysis of the diel cycle in *C. subtropica* confirmed: i) upregulation of photosynthesis and glycogen biosynthesis genes during the day and ii) upregulation at night of nitrogen fixation genes (*nif*), glycogen degradation and pathways that consume glucose, i.e. glycolysis, the oxidative pentose phosphate (OPP) pathway, and the tricarboxylic acid (TCA) cycle (34). Overall, these observations support a model whereby glucose produced by photosynthesis is stored as glycogen, which is catabolized at night to provide ATP and reducing equivalents for nitrogenase activity.

Because the diazoplast no longer performs photosynthesis, its nitrogenase is fueled by the products of host photosynthesis, i.e. photosynthate. To investigate the metabolic coupling between host photosynthesis and diazoplast nitrogenase activity, ARAs were performed under light, dark, and light + an inhibitor of photosystem II, 3-(3,4-dichlorophenyl)-1,1-dimethylurea (DCMU). During the day, inhibition of the host photosynthetic activity by dark condition or DCMU treatment reduced nitrogenase activity by 60% (Fig. 3*A*). At night, host photosynthetic activity by addition of light provided little to no benefit for nitrogenase activity (Fig. 3*A*). These results indicate that photosynthates are directly shuttled from the host chloroplast to the diazoplast during the day, while nighttime nitrogenase activity relies on carbohydrate storage. The new ability of daytime nitrogen fixation, concurrent with photosynthesis, permits a direct use of host photosynthate without reliance on carbohydrate storage. Moreover, in both conditions, blocking oxygen production with DCMU did not increase nitrogenase activity, indicating a mechanism to prevent inactivation of nitrogenase by the oxygen produced during host photosynthesis.

**Figure 3.**
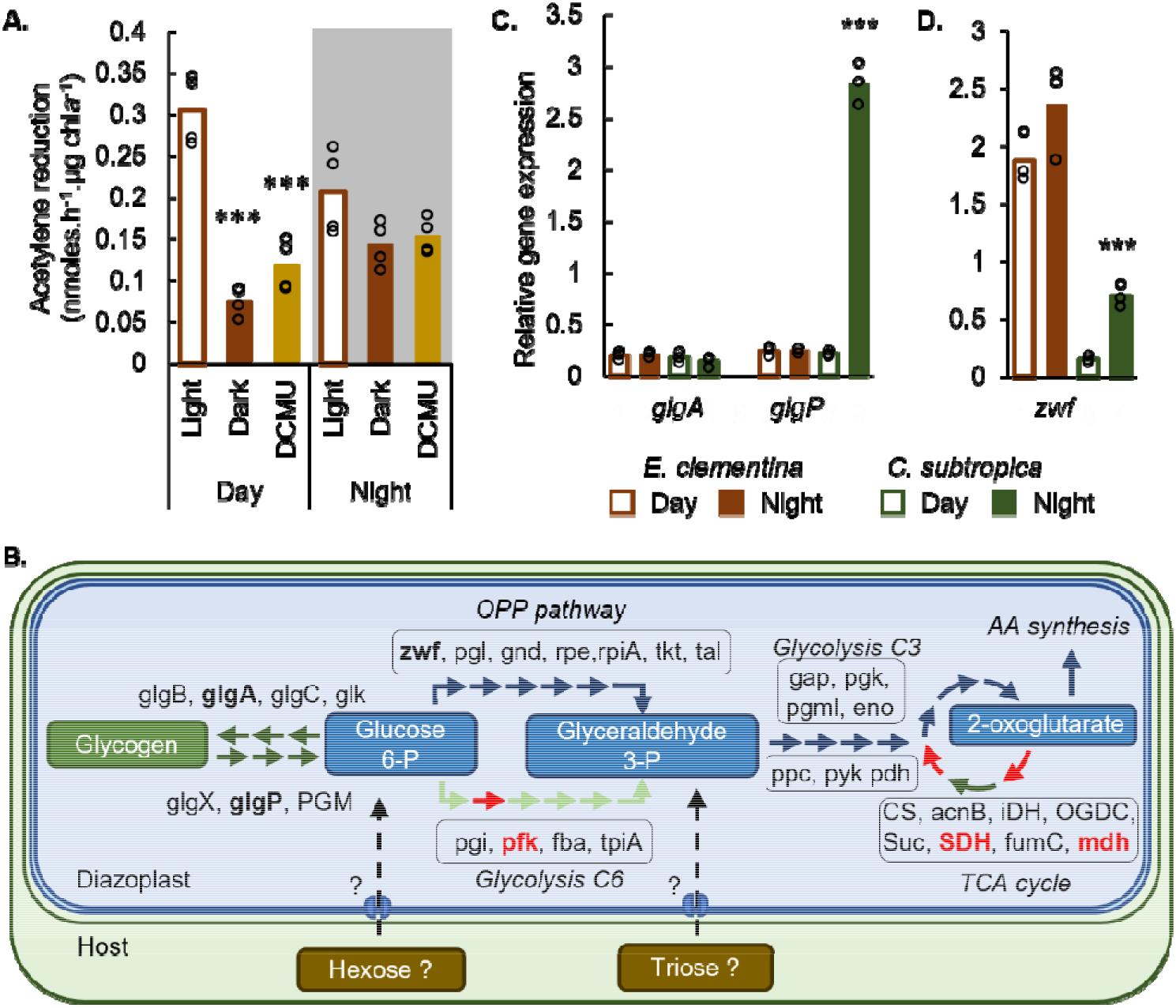
Carbohydrate metabolism associated with nitrogen fixation activity in the diazoplast. (A) Nitrogenase activity assessed by acetylene reduction assay at day and nighttime points in *E. clementina,* comparing photosynthetic conditions (light) and non-photosynthetic conditions (dark or DCMU treatment). (B) Annotated carbon metabolism pathways in diazoplast, showing proteins only present in *C. subtropica* and absent in diazoplast’s genome (red). See Table S1 for corresponding gene ID numbers. (C-D) Relative gene expression of representative genes in glycogen biosynthesis (*glgA*), glycogen degradation (*glgP*), and OPP (*zwf*) pathways in *E. clementina* (brown) and *C. subtropica* (green) during early day (open bars) and early night (solid bars). Mean values are plotted with individual biological replicates indicated by data points; *p*-values are relative to control condition.

To determine the source of energy fueling nighttime nitrogen fixation in the diazoplast, we assessed its glycogen metabolism. Despite gene reduction in glycogen metabolism pathways, the diazoplast genome encodes at least one copy of each gene required for glycogen biosynthesis and degradation, suggesting these pathways are functional (Fig. 3*B*, Table S2). Indeed, glycogen is common even in non-diazotrophic cyanobacteria as a storage for excess photosynthate produced during the day which is then catabolized at night (35). If the diazoplast accumulates glycogen as an intermediate storage form of host photosynthate to fuel nighttime nitrogen fixation activity, glycogen biosynthesis and degradation pathways are expected to be regulated in a diel pattern. Therefore, we measured the daytime and nighttime expression level of genes related to glycogen biosynthesis (glycogen synthase, glgA) and glycogen degradation (glycogen phosphorylase, glgP) in the *E. clementina* diazoplast and *C. subtropica* by RT-qPCR. Expression levels of *glgA* did not show significant variation between day and night timepoints in either *E. clementina* or *C. subtropica* (Fig. 3*C*). The expression level of *glgP* was upregulated by over 10- fold at night compared to the day in *C. subtropica*, consistent with the catabolism of stored glycogen to provide energy for nighttime nitrogen fixation in this organism. In contrast, no diel variation of *glgP* was seen in the diazoplast. In fact, *glgP* expression was consistently low in the diazoplast and comparable to daytime *glgP* expression levels in *C. subtropica* when the glycogen degradation pathway is effectively inactive (Fig. 3*C*). Overall, the low expression level and loss of diel regulation of the glycogen degradation enzyme, *glgP*, indicates that glycogen is not the source of energy for nitrogenase in the diazoplast, even during nighttime. Consistent with our results, glycogen granules were not detected in previous electron microscopy of diazoplasts from *Epithemia* spp. (17, 36). Since the diazoplast genome does not encode other pathways for carbohydrate storage, nighttime nitrogen fixation in the diazoplast most likely requires mobilization of host storage carbohydrates. Taken together, carbohydrate utilization in the diazoplast demonstrates tight coupling between host and diazoplast metabolism beyond that expected to support daytime nitrogen fixation.

### Carbohydrate catabolism is dependent on the OPP pathway in the diazoplast

After examining the potential sources of carbohydrate for daytime and nighttime nitrogen fixation in the diazoplast, we turned our attention to catabolism in the diazoplast to convert host-derived carbohydrates into ATP and reducing equivalents. *C. subtropica* has been shown to rely on glycolysis, the OPP pathway, and the TCA cycle for carbon catabolism (34). However, of these, the OPP pathway is the only complete pathway in diazoplasts, since many of the TCA cycle genes and the gene encoding the “gatekeeper” enzyme of glycolysis, phosphofructokinase (PFK), are missing from diazoplast genomes of *E. clementina*, *Epithemia turgida*, and *Rhopalodia gibberul*a (20) (Fig. 3*B* Table S2). In heterocysts, specialized nitrogen-fixing cells found in filamentous cyanobacteria, nitrogen fixation activity was shown to be highly dependent on glucose-6-phosphate dehydrogenase (G6PD), a key enzyme of the OPP pathway (37, 38). To investigate the importance of the OPP pathway in diazoplast metabolism, we measured the expression of *zwf* which encodes G6PD in the diazoplast compared with *C. subtropica*. The expression level of *zwf* in *E. clementina* was 2.5-3x higher than that of *C. subtropica* under nitrogen-fixing conditions, despite comparable *nifK* expression (Fig. 3*D* and 2*D*). Consistent with the loss of temporal regulation of nitrogen fixation, *zwf* expression did not exhibit diel variation in the diazoplast. Combined with the loss of key enzymes in glycolysis and TCA pathways, the moderate increase of *zwf* expression suggests that OPP is the main pathway for carbohydrate catabolism in the diazoplast (Fig. 3*B*). In heterocysts, the dependence on the OPP was proposed to maximize production of reducing equivalents for oxidative phosphorylation that both generates ATP and consumes oxygen. High levels of oxidative phosphorylation may therefore be important for protecting nitrogenase from inactivation by oxygen produced during photosynthesis. In addition, because G6PD utilizes glucose-6-phosphate as its substrate, the dependence on the OPP pathway in the diazoplast indicates that at least one host-derived carbohydrate is a hexose.

### The diazoplast responds to exogenous ammonia but paradoxically represses ammonium uptake

Diazotrophs show feedback regulation repressing nitrogenase activity when exogenous nitrogen, either in the form of ammonia or nitrate, is available. The utilization of these nitrogen sources is dependent on several gene products (Fig. 4*A*; Table S2). Ammonia can diffuse across membranes in its neutral form or be imported in its protonated form via an ammonium transporter (Amt). Intracellular ammonia is then assimilated into amino acids by glutamine synthetase (GS) and glutamine oxoglutarate aminotransferase (GOGAT). Utilization of nitrate is dependent on uptake via an ABC transporter followed by intracellular conversion to ammonia by nitrate and nitrite reductases. Consistent with an ancestral ability to utilize ammonia and nitrate, these pathways are conserved in both *C. subtropica* and a related cyanobacterium, *Rippkaea orientalis* (*Cyanothece* sp*. PCC8801*). In contrast, the genes for ammonia uptake and assimilation are retained in the diazoplast genomes from *E. clementina*, *E. turgida*, and *R. gibberul*a but those encoding the nitrate ABC transporter, nitrate reductase, and nitrite reductase are missing (Table S2).

**Figure 4.**
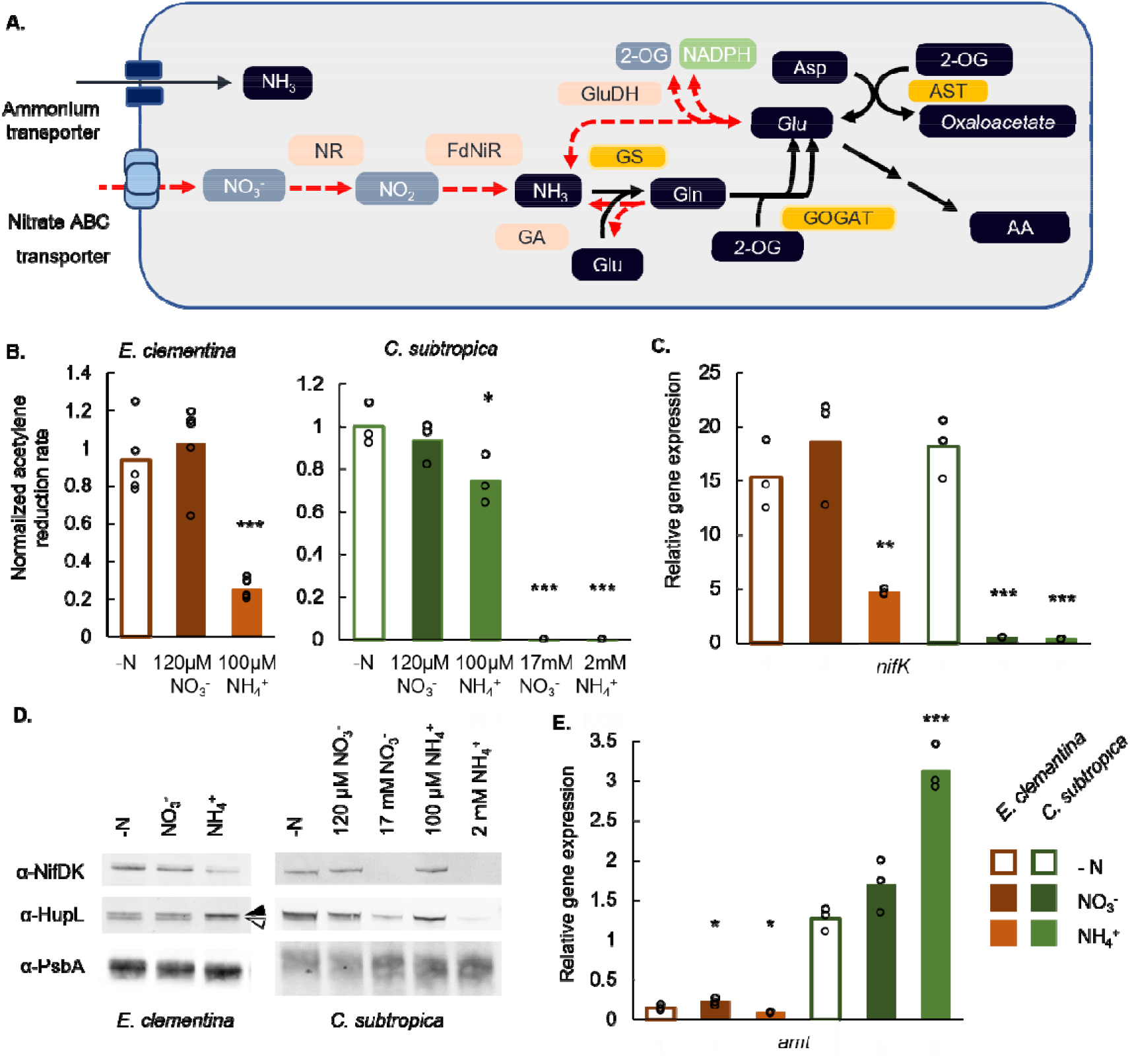
Effects of nitrogen repletion on nitrogen fixation. (A) Annotated nitrogen utilization pathways showing genes absent in the diazoplast but present in *C. subtropica* (red). See Table S1 for corresponding gene ID numbers. (B) Relative nitrogenase activity assessed by acetylene reduction assay upon repletion with nitrate or ammonium. Data were normalized by the mean of N-free samples. Measurements were collected at 2 hours into the day for *E. clementina* and 4 hours into the night for *C. subtropica*. (C) Relative expression of *nifK* in corresponding samples from B. For *C. Subtropica*, only samples in high concentrations of nitrate and ammonium were measured. (D) Immunoblots for NifDK, HupL, and PsbA proteins in corresponding samples from B. The black arrow points to the apo-HupL and the white arrow points to the mature HupL protein. (E) Relative expression of *amt* in corresponding samples from B. Mean values are plotted with individual biological replicates indicated by data points; *p*-values are relative to -N control condition.

Although *C. subtropica* is known to grow on nitrate or ammonium, its response to nitrogen repletion has not been documented (12, 29). We measured nitrogenase activity in *C. subtropica* in the presence of exogenous ammonium and nitrate at low (µM) and high (mM) concentrations. Neither 100 µM ammonium nor 120 µM nitrate strongly suppressed nitrogenase activity. However, higher mM concentrations fully repressed nitrogenase activity (Fig. 4*B*). This repression was reflected in loss of *nifK* expression by RT-qPCR (Fig. 4*C*) and of NifDK and HupL proteins by immunoblot (Fig. 4*D*). In *E. clementina*, repression of nitrogenase activity was observed at 100µM ammonium but unaffected by 120 µM nitrate (Fig. 4*B*). Ammonium repletion caused a 70% decrease in *nif* gene expression with a 2x decrease in NifDK protein abundance as well as a decrease in HupL abundance, which accumulated as a higher molecular weight apoprotein (Fig. 4*C* and *D*). These effects on nitrogen fixation were not observed with nitrate repletion, confirming the loss of nitrate utilization in the diazoplast. Consistent with a restricted ability to utilize nitrogen sources, diazoplast genomes are also missing functional genes for glutaminase and glutamate dehydrogenase required to catabolize amino acids for ammonia (Fig. 4*A*, Table S2). Curiously, the diazoplast was more sensitive to ammonium repletion, repressing nitrogenase activity at a lower ammonium concentration than *C. subtropica*.

Along with repression of nitrogen fixation activity, nitrogen repletion typically upregulates uptake and assimilation pathways to maximize utilization of exogenous nitrogen sources. Indeed, we observed that ammonium repletion increased *amt* expression by over 2-fold in *C. subtropica* (Fig. 4*E*). In the absence of exogenous nitrogen sources, the baseline expression of *amt* was 10- fold lower in the diazoplast than in *C. subtropica*. In contrast to upregulation observed in *C. subtropica*, ammonium repletion further decreased *amt* expression in the diazoplast (Fig. 4*E*). Altogether, the increased sensitivity to ammonium repressing nitrogen fixation combined with a diminished ability to utilize nitrogen sources in the diazoplast may optimize the host cell’s nitrogen resources.

#### Conserved regulatory genes show altered responses to exogenous ammonium

Considering the significant integration of diazoplast and host metabolism, two models for regulation of nitrogen metabolism could account for our observations: either the diazoplast retains cyanobacterial gene regulatory networks and senses its metabolic status (i.e. ammonia, carbon) for its response; or, regulatory genes are transferred to the host diatom implying a direct control of nitrogen metabolism in the diazoplast. Several transcription factors involved in nitrogen metabolism have been identified in cyanobacteria (Fig. 5*A*, Table S2): NtcA is a global regulator of nitrogen metabolism conserved in both diazotrophic and non-diazotrophic cyanobacteria (39–42). NtcA is activated by binding 2-oxoglutarate and PipX, which in turn is regulated by PII protein dependent on ammonia levels, effectively integrating information about the C:N balance of the cell (43–50). An important target of NtcA in diazotrophic cyanobacteria is CnfR (originally PatB), the master regulator of *nif* genes (51–53). This NtcA-CnfR regulatory network was identified in *C. subtropica* and in the diazoplast genome (Table S2). However, while the NtcA recognition sequence (TGTA-N_8_-TACA) can be identified upstream of *C. subtropica CnfR* (-449 bp to -434 bp) (40, 53), it is absent from the promoter region of *CnfR* in the diazoplast. To determine whether the NtcA-CnfR cascade still regulates the diazoplast *nif* cluster, we measured the expression of these key transcription factors in nitrogen-depleted and -repleted conditions. The relative expression of *ntcA* did not vary with ammonia repletion in either *C. subtropica* or diazoplast and had similar level of expression between the two, consistent with its being post-translationally regulated by metabolite levels and sensor proteins (Fig. 5*B*). *cnfR* expression showed a clear 58% decrease upon ammonium repletion in the diazoplast, with a similar 77% decrease upon nitrate or ammonium repletion in *C. subtropica* (Fig. 5*C*). The conserved responses in nitrogen fixation activity and its gene regulation between *C. subtropica* and the diazoplast suggest that the NtcA-CnfR regulation of the *nif* cluster has been retained in the diazoplast to activate nitrogen fixation activity in N-depleted conditions.

**Figure 5.**
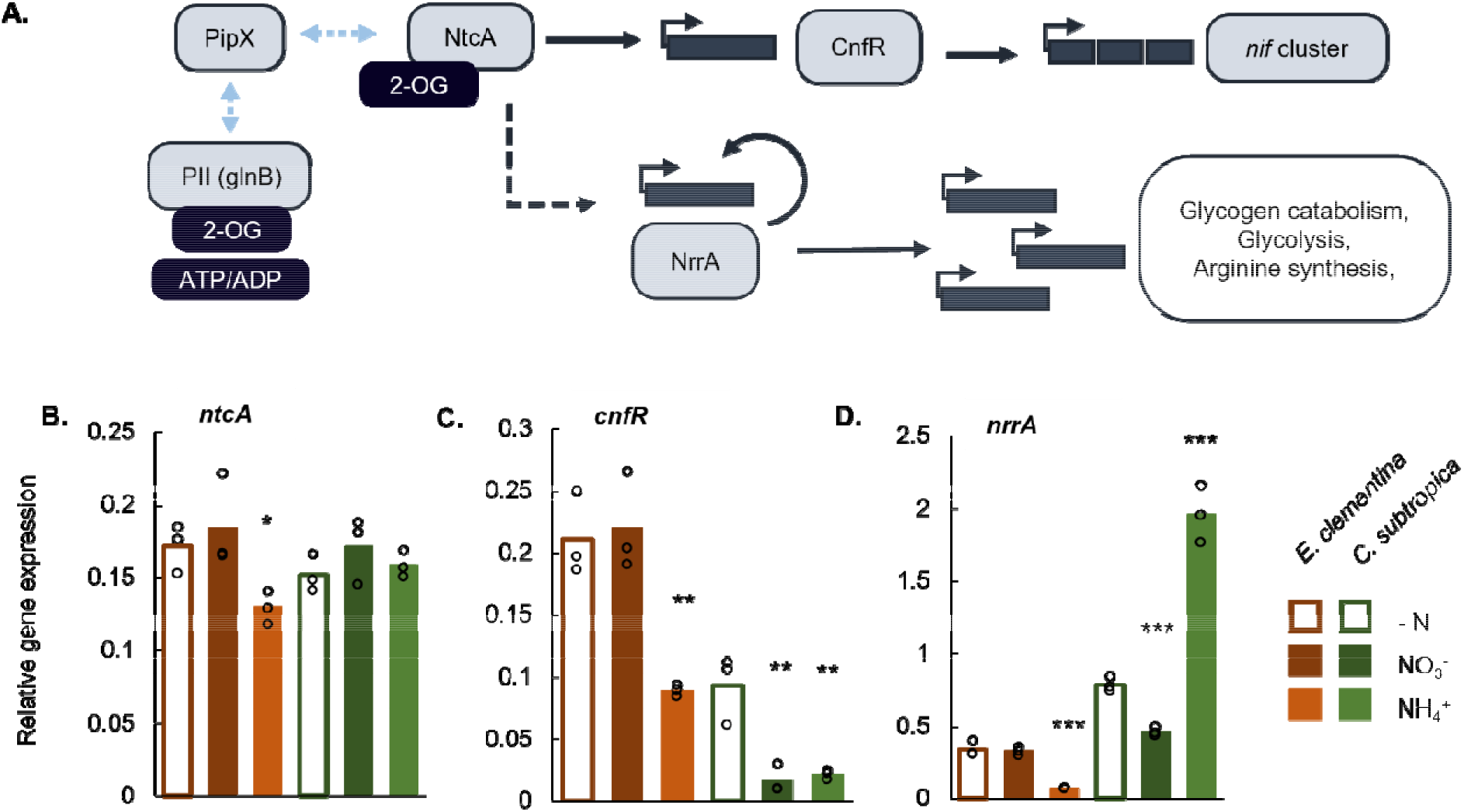
Effect of nitrogen repletion on regulatory genes. (A) Gene regulation for nitrogen fixation, nitrogen assimilation, and associated carbon metabolism in cyanobacteria. See Table S1 for corresponding gene ID. (B-E) Relative gene expression of (B) *ntcA*, (C) *cnfR*, (D) *nrrA*. Samples were taken at 2 hours into the day for *E. clementina* and 4 hours into the night using only high concentrations of nitrate and ammonium for *C. subtropica*. Mean values are plotted with individual biological replicates indicated by data points; *p*-values are relative to -N control condition.

NtcA also regulates the nitrogen-regulated response regulator, NrrA, a transcription factor of the OmpR family known to coordinates nitrogen and carbon reserves (54). A *nrrA* homolog was identified in *C. subtropica* and the diazoplast genome (Table S2). Of the transcription factors that we tested, NrrA gave the most unexpected response: upon ammonia repletion, *nrrA* was upregulated in *C. subtropica* but downregulated in *E. clementina* (Fig. 5*D*). Unfortunately, regulation of nitrogen metabolism has not been investigated in *C. subtropica,* and NrrA has diverse gene targets in cyanobacteria. Consequently, the downstream effects resulting from the altered *nrrA* response in the diazoplast is unclear. An *in silico* study, conducted in 15 different cyanobacteria, characterized target sequences of NrrA and experimentally validated them in *Synechocystis* sp. PCC 6803 (55). The proposed NrrA regulon in *C. subtropica* and *R. orientalis* included genes involved in glycogen catabolism, central carbon metabolism, arginine biosynthesis, and protein degradation. We manually screened the promoter region of the proposed targets and confirmed the presence of a potential NrrA binding sequence in the promoter region of *glgP*, *argG*, and *nrrA* itself in the diazoplast genomes of all *Epithemia* spp. (Fig. S3). The conservation of the binding sequences suggests that the NrrA regulon is conserved between *C. subtropica* and the diazoplast, but its response to exogenous nitrogen is altered upstream of *nrrA*. Once again, the paradoxical response of *nrrA* in the diazoplast compared to *C. subtropica* indicates key adaptations in nitrogen assimilation and storage that accompanied the endosymbiotic transition.

## Discussion

Metabolic exchange is often the underlying currency of symbiotic interactions and a strong driver of evolution. Our results demonstrate that the acquisition of nitrogen fixation by *Epithemia* diatoms has led to global metabolic rearrangements in the diazoplast and a level of metabolic integration not previously seen in other diazotrophic symbioses. Whereas *Crocosphaera* cyanobacteria, quintessential generalists, are free-living phototrophs that perform nitrogen fixation exclusively at night, the diazoplast has evolved into a nitrogen-fixing specialist and obligate heterotroph endosymbiont. In reviewing our key findings below, we discuss potential mechanisms for these metabolic changes and how these changes may be tailored to support nitrogenase activity.

***Mechanisms enabling daytime nitrogen fixation*.** The first notable change we observed was daytime nitrogen fixation activity in the diazoplast. We observed continuous nitrogen fixation activity in *E. clementina*, while recently discovered marine *Epithemia* species, *E. pelagica* and *E. catenate*, showed nitrogen fixation activity over the daytime with a hiatus of activity at night, a reversed day-night pattern compared to *C. subtropica* (13). This lax temporal regulation is in stark contrast to *C. subtropica*, in which 30% of genes, mostly associated with photosynthesis and nitrogen fixation, show circadian expression (compared to just 2-7% of genes in the non-diazotrophic cyanobacterium, *Synechocystis* 6803) (34, 56–58). Cyanobacterial circadian clock relies on the 3 *kia* genes and *C. subtropica* has 2 copies of each gene while the diazoplast retained 1 copy of each (Table S2) (59). It is not clear how the loss of redundancy would affect circadian rhythm. Alternatively, the loss of redox signals resulting from photosynthetic activity that serve as environmental cues to synchronize light and dark periods may weaken the circadian cycle. Finally, we hypothesize that the incomplete TCA cycle in the diazoplast may lead to accumulation of 2-oxoglutarate, which is both a substrate for GS-GOGAT-mediated ammonia assimilation and a signaling metabolite that binds to NtcA and PII (Fig. 4*A*). The accumulation of 2-oxoglutarate would signal a high C:N ratio in the diazoplast, a metabolic state that activates nitrogen fixation and may contribute to the temporal deregulation of nitrogen fixation. Regardless of the mechanism, the consistent shift toward daytime nitrogen fixation in all diazoplasts indicates a benefit of direct usage of photosynthate to fuel nitrogen fixation. The cyanobacterium *Trichodesmium*, which also performs nitrogen fixation concurrent with oxygenic photosynthesis, has been proposed to benefit from the absence of glycogen buildup during the day to maintain buoyancy in the water column (60, 61). In *Trichodesmium*, the mechanism by which nitrogenase is protected from the oxygen produced by photosynthesis is not known.

### What host-provided carbohydrate(s) is utilized by the diazoplast?

We observed that daytime nitrogen fixation was dependent on photosynthetic activity while nighttime nitrogen fixation was no longer associated with glycogen storage and degradation, suggesting the diazoplast relies entirely on carbohydrates imported from the host. Though photosynthetic, some cyanobacteria have been observed to grow mixotrophically on exogenous carbons, usually glucose, fructose, and/or sucrose. Two transporters, GlcP and FrtABC, have been associated with glucose and fructose uptake respectively, in cyanobacteria. However, orthologs of these transporters are not detected in *C. subtropica* nor diazoplast genomes. Similarly, utilization of the disaccharide sucrose requires an invertase to hydrolyze it into glucose and fructose, but no invertase has been detected in *C. subtropica* or diazoplast genomes. Instead, *C. subtropica* shows robust growth on glycerol as a carbon source and utilizes glucose after a prolonged period of adaptation (11, 12). The transporters that allow *C. subtropica* to readily uptake glycerol and adapt to glucose uptake are unknown, though potential genes annotated as sugar ABC transporters are identified in the genomes of *C. subtropica R. orientalis*, and three diazoplasts. More detailed investigations of mixotrophic growth in *C. subtropica* will be needed to determine whether pre-existing transporter gene(s) were utilized to access host carbohydrates. An alternative model is that the diazoplast acquired the ability to uptake host carbohydrate(s), since we detected transporter genes present in all diazoplast genomes that were absent from free-living *Crocosphaera* genomes. Further investigation is needed, but these could be exciting transporter candidates. We also considered the possibility that, instead of sugar compounds, ATP and NADPH could be directly imported into the diazoplast. However, none of the transporters known in mitochondria or chloroplasts were identified in diazoplast genomes, and the observed high expression level of *zwf* suggests the import of hexose from the host that is metabolized through the OPP pathway.

### The OPP pathway and oxidative phosphorylation provide reducing equivalents, ATP, and protection from oxygen for nitrogenase

.The third change we noted was an incomplete set of glycolysis and TCA cycle genes and likely dependence on an upregulated OPP pathway for carbohydrate catabolism. While multiple enzymes in the TCA cycle were missing such that it was unlikely to be reconstituted, only PFK, the enzyme responsible for the first committed step of glycolysis, is absent from the diazoplast genome compared to C. subtropica. Loss of PFK is relatively common in cyanobacteria; however, we also considered the possibility that the PFK gene may have transferred to the diatom nucleus and may be imported. Compared to glycolysis, the OPP pathway favors production of reducing equivalents at the expense of ATP production. However, these reducing equivalents can then enter the respiratory electron transport chain. Electron micrographs of the E. gibba diazoplast show the presence of thylakoid membranes (17, 36). Unlike plant chloroplasts, both photosynthetic and respiratory electron transport complexes are found on thylakoid membranes in cyanobacteria. Therefore, in the absence of photosynthesis, diazoplast thylakoids likely were retained for respiratory electron transport and oxidative phosphorylation. We propose that, in the diazoplast, metabolism of sugars via the OPP pathway and the latter “pay-off” phase of glycolysis generates reducing equivalents for oxidative phosphorylation, which in turn not only provides the relatively large equivalents of ATP needed for nitrogenase but also consumes oxygen as the terminal electron acceptor maintaining an anoxic microenvironment for nitrogenase activity. Consistent with this model, heterocysts are dependent on the OPP pathway for their nitrogenase activity, and genes related to oxidative phosphorylation are upregulated at the transition from day to night in C. subtropica (34, 37, 38). There may be other mechanisms for scavenging oxygen in the diazoplast that we are continuing to explore.

### A model for nitrogen transfer to the host

Finally, we demonstrated that while the negative feedback regulation of nitrogen fixation to ammonium was intact, and even sensitized, the same genetic regulation and environmental stimulus (i.e., ammonium repletion) resulted in paradoxical nitrogen assimilation responses and general suppression of the ammonia transporter, Amt. These results were puzzling at first but appear consistent with changes observed in diazotroph Rhizobium bacteria that colonize the root nodule of legumes. The Rhizobium amt gene is downregulated during its differentiation into nitrogen-fixing bacteroids in root nodules (62). Moreover, ectopic expression of amt disrupts bacteroid differentiation and the development of symbiotic nodules (63). Based on these findings, a model of nitrogen transfer in the legume nodule was proposed: ammonia produced by nitrogen-fixing bacteroids diffuses into the symbiosome space, an endosymbiotic compartment containing the bacteroids. Acidification of this space mediated by V-type H^+^-ATPases traps ammonia in its protonated ammonium form at high concentrations (64). The downregulation of the bacteroid amt has been proposed to prevent ammonium transport back into the endosymbiotic bacteria; instead, ammonium is transported across the symbiosome membrane into the plant cell. In analogy to the root cell symbiosome, ammonium may also accumulate in the endosymbiotic compartment of the diazoplast. This ammonium accumulation around the diazoplast, is consistent with the increased sensitivity of the diazoplast to exogenous ammonium and the general suppression of amt expression compared to C. subtropica. As a result, the diazoplast is maintained in a state of nitrogen deficiency. Combined with the increased accumulation of 2-oxoglutarate (discussed above), the high C:N balance may positively regulate the expression of nif genes. We hypothesize that this mechanism has evolved to ensure nitrogen fixation activity to supply the host diatom’s needs.

### Implications for engineering nitrogen-fixing crop plants

Crop plants, particularly cereals, with the ability to fix nitrogen are the holy grail for sustainable agriculture. However, so far, attempts to transfer the genes for biological nitrogen fixation from bacteria into plants have met many challenges, including the large number of genes required, the complexity of nitrogenase assembly, the high cellular energy requirements, and exquisite sensitivity to oxygen (5, 6, 65). Ironically, the current strategy relies on targeting nitrogenase to the mitochondria where high respiratory rates in the compartment provides energy and protects it from inactivation (66–69). Although, where the diazoplast is on the evolutionary path between endosymbiont and organelle is unknown, it serves as a ready-made nitrogen-fixing factory, expressing all the genes required, properly regulated, and housed inside a protected membrane compartment. Given the advantages of nitrogen fixation capability, the absence of such an organelle is a mystery but may be an accident of evolutionary timing or a result of incomplete documentation of eukaryotic diversity (70, 71). Regardless of the status of diazotroph endosymbiosis in nature, increasing related knowledge and new genetic tools open the opportunity to revisit this strategy in the laboratory. We propose that *Epithemia* diazoplasts could serve as a blueprint for engineering nitrogen-fixing organelles in plants. The metabolic adaptations we have observed in the diazoplast, for example, offers several options for genetic engineering of a suitable bacterial partner. We are currently unaware of the extent of the host contribution to the endosymbiotic interaction, whether solely in the form of metabolites or also via imported gene products. However, analysis of *Epithemia* genomes is underway in our laboratory and will provide insight into gene transfers and imported gene products, if any. Altogether, our investigations of this native endosymbiotic interaction will be a powerful tool for bioengineering.

## Materials and Methods

### Strain isolation and cultivation, and microscopy

Environmental samples originated from Gazos Creek near Butano State Park, CA, USA and were taken during fall 2020. *E. clementina* was isolated by successive dilution in a nitrogen-depleted media, Csi-N (Table S3), under 10 µmole photon.m^-2^.s^-1^ of white light at 20°C as previously described (26) and have been since then been maintained in the same condition. After subculturing for 8 weeks, we obtained many monoalgal isolates, some containing filamentous cyanobacteria and some containing *Epithemia* diatoms. Cultures, used in experiment, were started from single cell isolate of *E. clementina*. Cultures were monoalgal but not axenic, and cells formed a biofilm on the surface of the flasks. The Csi-N media was complemented with reduced nitrogen by addition of Ca(N0_3_)_2_, 4 H_2_0 at a final concentration of 60 µM or (NH_4_)_2_HPO_4_ at a final concentration 100 µM as described in (17). Conditions were changed 3 days prior to the measurements (Acetylene reduction assays and mRNA extraction).

*Crocosphaera suptropica* (*Cyanothece* ATCC51142) was kindly provided by Dr. Jonathan Zehr and maintained at 30°C under day-night cycle (12 h/12 h – 30 µmole photon.m^-2^.s^-1^/dark) in artificial sea water ASP2 (29) with constant bubbling of humidified ambient air. ASP2 media that contained no source of combined nitrogen, was complemented either as previously described for Csi media or with NaN0_3_ at a final concentration of 17 mM or ammonium (NH_4_)_2_HPO_4_ at a final concentration of 2 mM (12). Conditions were changed 3 days prior to the measurements.

Observation of diazoplasts by DNA stain was performed after fixation for 10 min with 4% paraformaldehyde followed by incubation for 30 min with 1µg/mL DAPI. Fluorescence microscopy was performed with a CY5 and TRITC filter for chlorophyll and phycoerythrin respectively.

## Sequencing of the diazoplast genome

*Epithemia clementina* gDNA was extracted using QUIAGEN Dneasy Plant Pro Kit [69206]. DNA fragments were prepped for Illumina sequencing with NEBext Ultra II FS DNA Library Prep Kit for Illumina [E7805S]. Sequencing was performed on an Illumina NextSeq 2000 P3 paired end 150bp. Sequencing reads were trimmed, quality filtered with fastp (72) (--qualified_quality_phred 20 –unqualified_percent_limit 20), assembled with SPAades on default settings (73) and quality was assessed with FastQC (74).

For Nanopore sequencing, gDNA was isolated with circulomics nanobind plant nuclei DNA kit. Short DNA was removed with circulomics short read eliminator kit. The remaining DNA was then library prepped for nanopore sequencing using the Genomic DNA by Ligation (SQK- LSK111 kit) and loaded onto an R 10.34 flow cell. Nanopore reads were basecalled with guppy and short (< 3000 bp) and low quality (Phred quality score < 10) reads were removed with nanofilt (75). Nanopore reads were assembled with Shasta (76), and contigs with one or more high quality (Expect value <= 10^-10^) BLAST hits to previously published diazoplast genomes (NCBI accessions NZ_AP012549.1, NZ_AP018341.1) (20, 21) were isolated to create a fragmented and incomplete draft assembly. Nanopore and 11riton11e reads were mapped to the draft assembly with minimap2 (77) and BWA-MEM (78) and mapped reads were re-assembled with unicycler (79) to produce a high-quality circular diazoplast genome. Genome quality was assessed with Merqury and quast (80, 81). Gene annotation of the diazoplast was performed with NCBI Prokaryotic Genome Annotation Pipeline.

### Phylogeny

Using the corresponding *Phaeodactylum tricornutum* or *Cyanothece* sp. ATCC 51142 gene sequences as queries, sequences of *Epithemia clementina* host genes for *psbC, rbcL* and 18S- rRNA and endosymbiont genes for *nifH* and 16S-rRNA were extracted from the Spades assembly by command-line BLAST (82, 83). Sequence lengths extracted are as follows: 1391bp of *rbcL*, 1076bp of *psbC*, 1622bp of 18S-rRNA, 761bp of *nifH,* and 1410bp of 16S-rRNA. A nucleotide BLAST against the NCBI nr/nt database was performed to ensure correct identity of these sequences. Sequences used for phylogeny are gathered in Table S4. For each gene, sequences were aligned using MAFFT v7.490 (mafft-linsi –adjustdirectionaccurately –maxiterate 1000) (84). Gaps and highly variable regions in the alignment were removed with trimAl v1.2rev59 (85) using the gappyout flag and inspected by eye to ensure proper alignment and trimming. Concatenated alignments and data partitions were generated using SequenceMatrix v1.9 (86). Phylogenetic trees were inferred using IQ-TREE 2 (87) with ModelFinder (88) automatic model selection and node support tested with 2000 iterations of rapid phylogenetic bootstraps.

### Metagenomic analysis and assembly

Metagenomic assembly was performed with flye (89) and contigs were polished 3x with racon (90). Eukaryotic contigs were removed with EukRep (91), and metagenomic binning was done with MetaBat2 (92). To investigate the possibility of nitrogen fixation by a free-living microbe, assembled contigs were searched against the UniProt reference proteomes database using Diamond (93) and results annotated as NifK (interpro IPR005976), NifD (interpro IPR005972), or NifH (interpro IPR005977) were isolated to identify putative nitrogen fixing microbes. To assess metagenomic sequence diversity, MAGs were classified using kraken2 (94) and metagenomic diversity was visualized with Krona (95).

### Nitrogenase activity assay

For acetylene reduction assays (ARA), *E. clementina* cultures were scraped off the flasks and centrifuged (1,500xg, 4 min). They were washed once in fresh Csi-N media and resuspended in the appropriate media. From that culture, 2.3 mL was placed in an autoclaved 10 mL glass vial and sealed with a breathable sealing film. Note that the volume of culture was set to 2.3 mL to leave a 10 mL headspace in the vial. Vials were placed in their respective growth conditions for 3 days before the assay. For *C. subtropica*, fresh cultures were grown for 3 days in the desired conditions as described previously and sampled for ARA at the time of the experiment.

Vials with 2.3mL of culture were sealed and injected with 1 mL of acetylene generated using calcium carbide and water. After one hour of incubation under culture conditions or desired conditions, the reaction was blocked by injection of 150 µL of 16% paraformaldehyde and kept at 4°C until further analysis. Ethylene, present in the 10 mL-head space, was quantified by injection of 1mL of the headspace on a gas chromatography coupled with a flame ionization detector, Shimadzu GC-8A1F. A Porapak N 80/100 mesh 6’ x 1/8” x 0.085” SSP/W column was used. The injector was set at 120°C and column at 80°C. Nitrogen was used as a carrier gas at 225kPa. Results were collected using a Shimadzu Chromatopac which performed automatic pic detection and quantification. A 1% ethylene standard gas was used to quantify the ethylene per signal detected. After gas analysis, cells were collected by centrifugation and chlorophyll was extracted with 100% ethanol for quantification (96).

### Protein extraction and immunodetection

Cells were scraped and harvested at different time points of the day-night cycle by centrifugation at 3,000xg for 2 min. Pellets were deep-frozen in liquid nitrogen and stored at -80°C until protein extraction. Pellets were resuspended in 200µL of lysis buffer (10mM HEPES, 10mM EDTA, 0.5% 12riton-100, 2mM DTT). Cells were homogenized with a mix of 1mm and 0.5mM glass beads at 3000 strokes per minute for 2 minutes in a bead beater. Lysate was centrifuged at 15,000g for 5 min, 4°C and only supernatant was kept. Six volumes of cold acetone (-20°C) were added. After 1h at -20°C, proteins were pelleted at 15,000 g for 10 minutes, 4°C. Supernatant was kept for chlorophyll concentration measurement (96). Pellet was washed with 80% cold acetone, gently dried and resuspended at a final concentration of 0.1 µg/mL of chlorophyll which correspond to 1 µg/mL of total protein content in 1X lithium dodecyl sulfate loading buffer with 100 mM DTT. Sample were denaturated at 70°C for 30 min, and vortexed periodically. Before loading, samples were pelleted at 15,000xg for 5 min. Precision Plus Protein™ All Blue Standards was used as a molecular ladder to assess protein molecular weight.

Proteins were separated by electrophoresis on NuPage Bis-Tris Gels, 4-12% Acrylamide using MES buffer. They were then transferred using Bio-rad Trans-blot Turbo transfer onto a nitrocellulose membrane. Membranes were blocked in LiCOR blocking buffer (0.1% Casein, 0.2x PBS, 0.01% sodium azide) for 1 hour at room temperature. The FeMo nitrogenase (NifDK subunits) was immunoblotted with polyclonal goat-raised antibody (1:500 dilution) kindly provided by Dr. Dennis Dean from Virgnia Tech, US. The large subunit of the hydrogenase (HupL) was immunoblotted with antibody (1:2500 dilution) raised in rabbit against amino acids 260 to 270 of HupL from *Anabaena* sp. PCC 7120 kindly provided by Dr. Paula Tamagnini from the Faculty of Sciences of University of Porto, Portugal. PsbA was used as an internal loading control; a 1:10,000 dilution was use of the rabbit antibody against the C-terminal of PsbA (AgriSera AB, Vanas, Sweden). Antibodies were diluted in 50% TBS-T / 50% LiCOR blocking buffer. After 2 hours with primary antibodies at room temperature, the membrane was washed 3 times with TBS-T and then incubated with LiCOR secondary antibodies (IRDye 800CW) for 1 hour (α-rabbit for PsbA and HupL and α-goat for NifDK). The membrane was rinsed twice with TBS-T and once with PBS and the blot was imaged using an infra-red LiCOR imager. Intensity of the signal was quantified using Image Studio Lite software v5.2. As FeMo nitrogenase and HupL have close molecular weight, the nitrogenase was first immunoblotted and the membrane was stripped with 0.2M NaOH for 5 minutes, washed with water twice and immunoblot was repeated for HupL.

### Gene expression

After 3 days of subculturing in the desired condition, cells were harvested at 1,500x*g* for 2 minutes and resuspended in Trizol Reagent (Invitrogen™, Carlsbad CA, USA) and then lysed by a combination of flash-freezing and sonication. mRNA was extracted using a Quiagen Rneasy® Plus Universal kit. mRNA concentration was assessed by nanodrop. Following extraction, RT-qPCR was performed with NEB Luna® Universal One-Step RT-qPCR using 125ng of mRNA per 10µL reaction as described in instructions. Amplification was performed and monitored in a 96- well plate using a StepOne^TM^ Real-Time PCR system according to the instruction of the Luna kit and was followed by a melting curve. Primers used are detailed in Table S5. Specification of the amplification was validated first by PCR on gDNA and confirmed after the RT-qPCR of mRNA from analysis of the melting curves. DNA gyrase subunit B and 30S SSU ribosomal protein S1p were used as housekeeping genes. Expression was normalized to weighted average of GyrB and 30S SSU expression.

**Fig S1:**
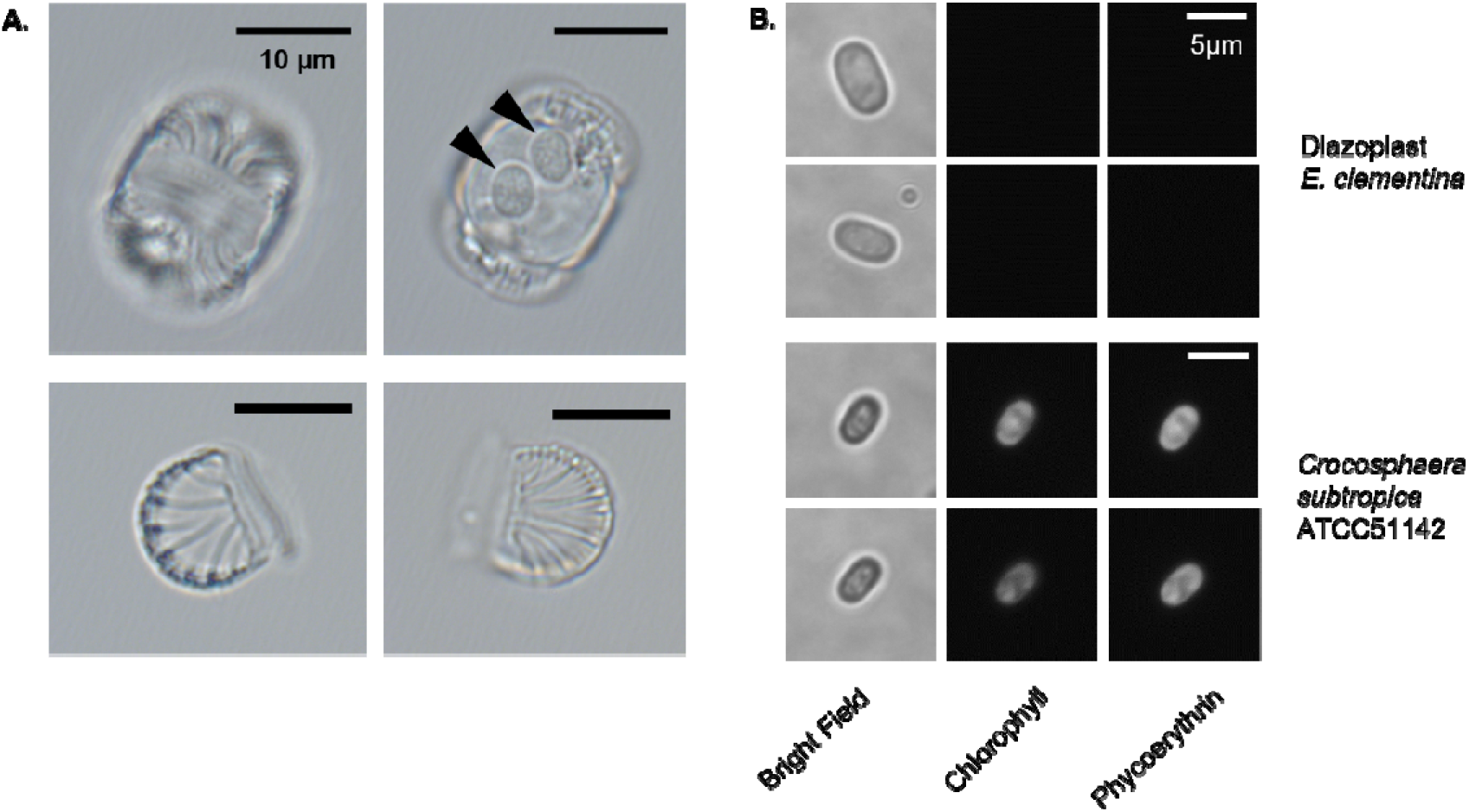
Supplemental micrographs of *E. clementina*. (A) Micrography of *Epithemia clementina* shell with diazoplasts (indicated by the black triangles) (B) Fluorescence micrography of *E. clementina* diazoplast (top) and *C. subtropica* (bottom) showing fluorescence of chlorophyll and phycoerythrin

**Fig. S2:**
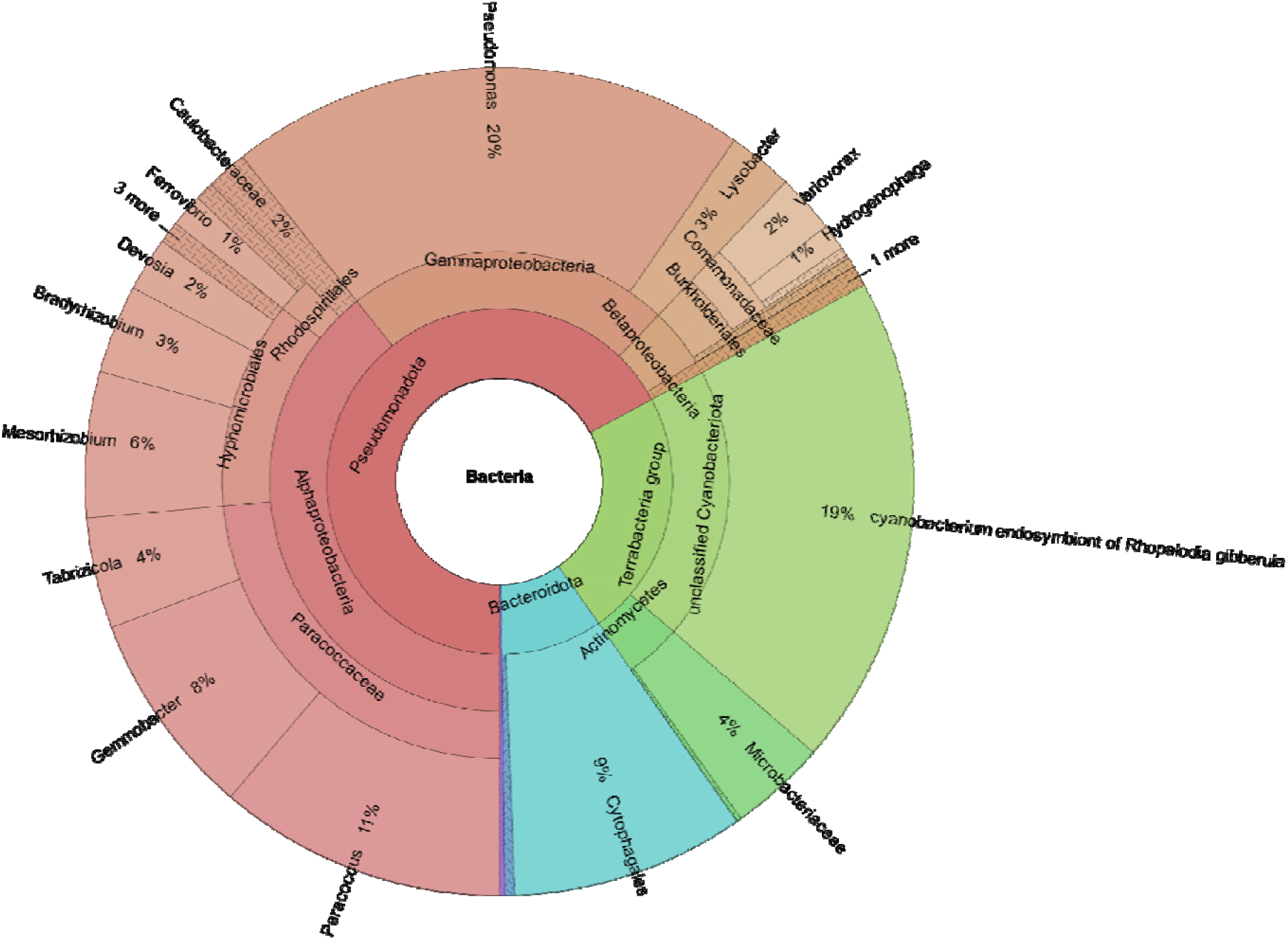
Diversity of the bacterial community. Krona visualization of kraken assignment of metagenome-assembled genomes. See Table S1 for detailed abundance.

**Fig. S3:**
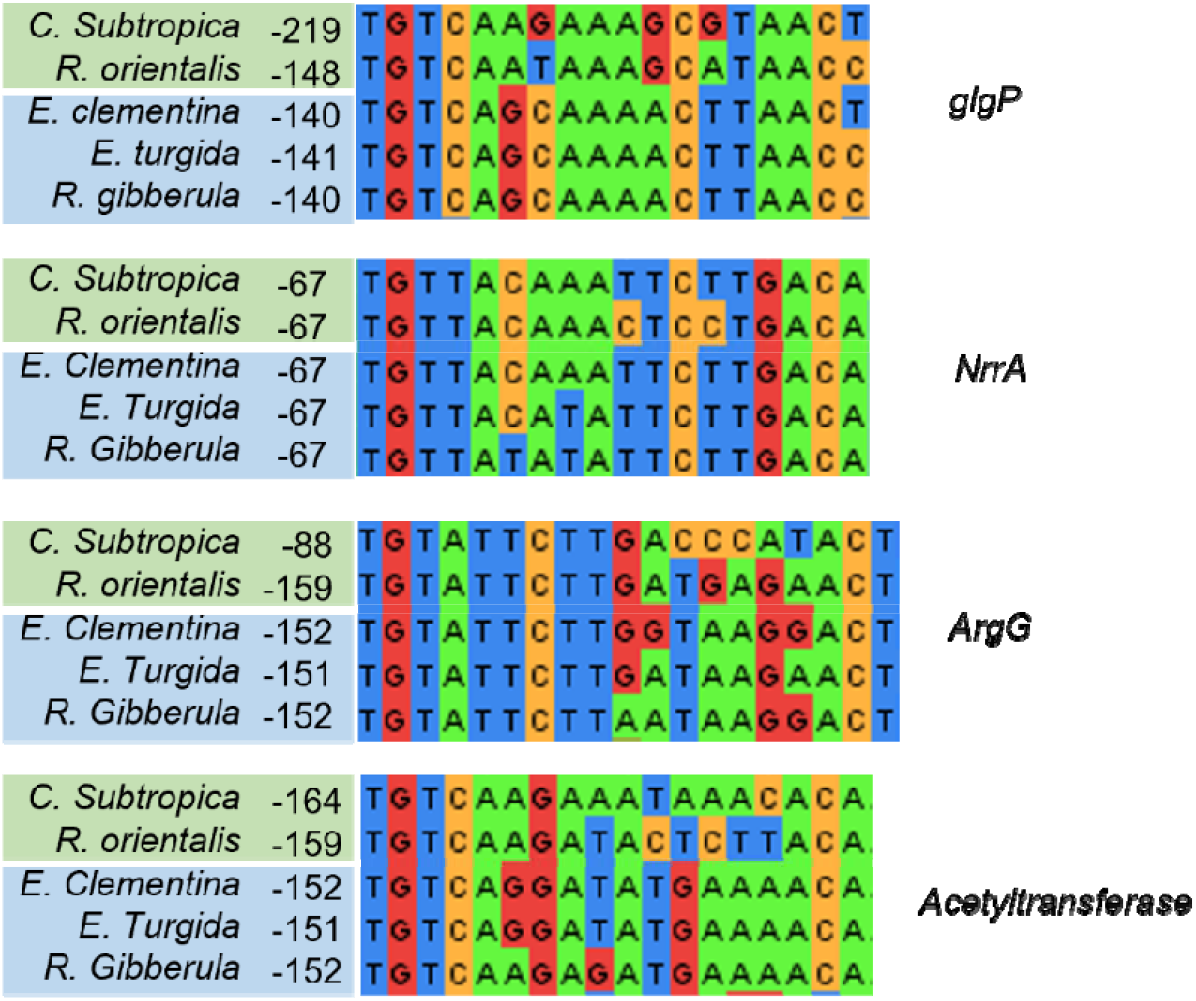
Regulon of NrrA in Cyanothece and diazoplast. Alignment of identified NrrA binding sequences in promoter region of *glgP*, *NrrA*, *ArgG* and an *Acetyltransferase* in two free-living Cyanothece (*C. subtropica* and *Rippkaea orientalis*) and 3 diazoplasts (*E. clementina*, *E. turgida*, *R. gibberula*). Number indicates the position from the start codon.

Table S1: MAGs analysis and taxonomic assignment.

Table S2: **NCBI gene IDs.** Genes present ID corresponding to protein involved in carbon metabolism, nitrogen metabolism, nitrogen regulation and circadian clock.

Table S4: Source data for phylogeny

Table S3: Media recipe for Csi-N.

| | Stock concentration<br>(M) | Final concentration<br>(μM) | Volume in 1L of<br>stock solution (mL) |
| --- | --- | --- | --- |
| CaCl<sub>2</sub> | 0.1 | 272 | 2.72 |
| K<sub>2</sub>HPO<sub>4</sub> | 0.1 | 57 | 0.57 |
| β-Na<sub>2</sub> glycerophosphate* | 0.1 | 163 | 1.63 |
| MgSO<sub>4</sub> | 0.1 | 162.3 | 1.623 |
| Na<sub>2</sub>SiO<sub>3</sub> 9H<sub>2</sub>O | 0.1 | 351.9 | 3.519 |
| Hepes | 1 | 10 mM | 10 |
| PIV Metals | - | - | 3 |
| B12 vitamin* | 1 mg/mL | 50 μg/L | 0.05 |
| Biotin* | 2.5 mg/mL | 50 μg/L | 0.02 |
| Thiamine HCl* | 1 mg/mL | 10 μg/L | 0.01 |

PVI Metals
|  | mg/100 mL | Concentration mM |
| --- | --- | --- |
| Na<sub>2</sub>EDTA 2H<sub>2</sub>O | 100 | 2.69 |
| FeCl<sub>3</sub> 6H<sub>2</sub>O | 19.6 | 0.73 |
| MnCl<sub>2</sub> 4H<sub>2</sub>O | 3.6 | 0.18 |
| ZnCl<sub>2</sub> | 1.04 | 0.076 |
| CoCl<sub>2</sub> 6H<sub>2</sub>O | 0.4 | 0.017 |
| Na<sub>2</sub>MoO<sub>4</sub> 2H<sub>2</sub>O | 0.25 | 0.010 |

Media was adjusted to pH 7.5 and autoclaved for 20 minutes.
Solutions marked with (\*) were filter-sterilized separately and added after autoclave.

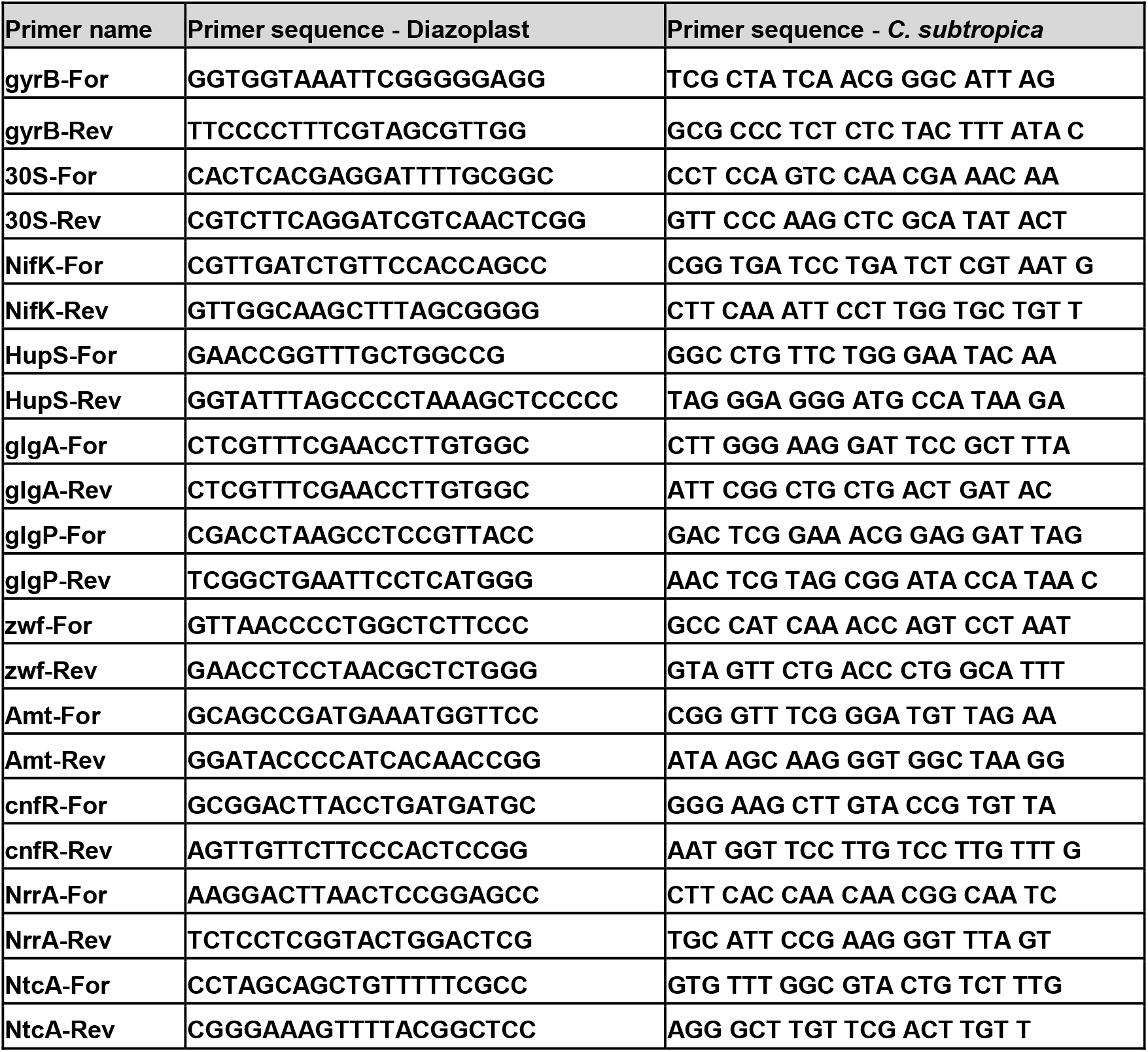
Table S5: List of primer used for RT-qPCR.

## Acknowledgments

We thank Dr. Jonathan Zehr, Dr. Devaki Bhaya, and Dr. Arthur Grossman for their feedback on the project and reviews on the manuscript. We are grateful for Dr. Jonathan Zehr, Dr. Sharon Long, Dr. Dennis Dean, and Dr. Paula Tamagnini to have shared material and equipment to make this work possible. E.Y. is a Chan Zuckerberg Biohub – San Francisco Investigator and supported by Burroughs Wellcome Fund. S.F. was partially funded by NIH training grant (T32GM007276)

## Notes

### Competing Interest Statement

The authors have declared no competing interest.

### Summary of Updates

Figure layout and improvement of the text.

